# Sex Bias in Iron Sequestration by Transferrin 1 Modulates Sexually-Dimorphic Infection Outcomes in *Drosophila melanogaster*

**DOI:** 10.1101/2025.08.07.669156

**Authors:** Alexandra Hrdina, Igor Iatsenko

## Abstract

Host sexual dimorphism in the outcome of infections is a ubiquitous phenomenon across taxa. However, the immunological differences between males and females and the mechanisms underlying them remain poorly characterized. Here, we used *Drosophila melanogaster* to test the hypothesis that sex differences in nutritional immunity, particularly iron sequestration, contribute to sexual dimorphism in infection outcome. Using the natural *Drosophila* pathogen *Providencia alcalifaciens*, which is controlled by host-mediated iron sequestration, we established an infection model in which males demonstrate increased resistance. Leveraging this model, we demonstrated that males exhibit higher basal and infection-induced expression levels of Transferrin 1-an iron transporter mediating iron sequestration during infection. Consequently, males contained lower iron levels in the hemolymph compared to females. Importantly, sex differences in iron content and in survival after *P. alcalifaciens* infection were abolished in *Tsf1* mutants. However, these mutants still exhibited sex differences in pathogen load. Thus, while Tsf1 mediates sexual dimorphism in iron sequestration and susceptibility to *P. alcalifaciens* infection, it also has an unexpected role in host tolerance to infection. Finally, we demonstrated that the Toll pathway mediates sex differences in *Tsf1* expression and susceptibility to infection. Altogether, our study demonstrates that Tsf1-mediated iron sequestration differs between male and female *D. melanogaster*, thereby identifying nutritional immunity as a determinant of sexual dimorphism in infection outcome.

## Introduction

Iron plays an essential role in numerous biochemical processes and is therefore vital for the survival of almost all life forms. It can be found as a cofactor in a wide variety of proteins, mediating processes such as oxygen transport, gene regulation and metabolism. The natural ability of iron to cycle between two oxidation states, Fe2+ and Fe3+ renders iron an important redox catalyst, which is highlighted by the fact that iron is the most abundant redox metal in biological systems (Andreini et al. 2008; Hood and Skaar 2012; Cassat and Skaar 2013). Nonetheless, iron levels need to be tightly controlled as free iron and reactive oxygen species (ROS) catalyze a Fenton reaction, generating hydroxyl radicals that damage cells (Cassat and Skaar 2013; Gomes et al. 2023). Subsequently, iron does not occur in its free form within the organism but is instead bound to iron-binding molecules and proteins, such as heme, transferrin and lactoferrin (Kaplan and Ward 2013; Cao et al. 2023).

The importance of iron in sustaining life raises issues at the interface between hosts and pathogens during infection. While hosts acquire iron via dietary intake or modifying the regulation of storage and use, pathogens rely on iron to be present in their immediate environment (Hood and Skaar 2012; Golonka et al. 2019). Host tissues present an iron-limited environment as the present iron is bound to iron-binding proteins and therefore unavailable to the pathogen. Furthermore, hosts will further restrict pathogen iron-access by sequestering iron from infected tissue. Immune responses, such as these, which involve transition metals, are generally termed nutritional immunity (Lopez and Skaar 2018; Hrdina and Iatsenko 2022; Murdoch and Skaar 2022). Conversely, pathogens have developed sophisticated strategies to overcome these defenses in order to successfully colonize. One prominent strategy is the production of siderophores, iron chelating molecules that have a higher affinity to iron than the iron-binding proteins produced by the host (Hider and Kong 2010; Golonka et al. 2019; Kramer et al. 2019; Helmann 2025).

*Drosophila melanogaster* is a widely appreciated model organism to study host-pathogen interactions due to its genetic amenability (Westlake et al. 2024). Curiously, nutritional immune responses have been understudied in the fruit fly (Missirlis 2021; Hrdina and Iatsenko 2022). The best investigated immune response involving transition metals in *Drosophila* is iron sequestration mediated by the iron transporter Transferrin-1 (Tsf1) (Xiao et al. 2019). Following infection, *Tsf1* expression has been shown to be induced via Toll- and IMD signaling and is responsible for removing iron from the hemolymph and shuttling it into fat body cells. Tsf1 deficient flies are more susceptible to multiple pathogens like *Pseudomonas* and *Providencia* bacteria as well as Mucorales fungi, highlighting the importance of iron in the defense against infection (Iatsenko et al. 2020; Shaka et al. 2022; Hrdina et al. 2024).

Biological sex has great influence on the physiology of an organism (Millington and Rideout 2018; Dähn and Wagner 2025). Sex differences are most obvious in anatomy and behavior but have also been reported in gene expression, epigenetics and hormone levels (Ratnu et al. 2017; Snell and Turner 2018; Dähn and Wagner 2025). Consequently, males and females respond differently to a variety of stressors, including infection and inflammation (Khan and Graze 2024; Rubinić et al. 2025). Even though sexual dimorphism has been reported in immunity (Klein and Flanagan 2016; Takahashi and Iwasaki 2021), research on this topic has been severely impacted missing reports of biological sex and a preference for unisexual studies within the field (Belmonte et al. 2020). Furthermore, the study of sexual dimorphism in immunity is made more complex by the fact that it is also dependent on the pathogen (D.F. Duneau et al. 2017; Belmonte et al. 2020). Hence, the mechanisms underlying sex differences in infection outcome remain poorly characterized.

It has previously been demonstrated, that male and female flies differ in the survival after pathogen challenge (Belmonte et al. 2020). Male flies were more resistant against infection with *Providencia alcalifaciens* and *Providencia rettgeri*, while females were less susceptible to *Staphylococcus aureus*. Moreover, it was shown that the regulation of the Toll pathway underlies sexual dimorphism and is more strongly induced in male flies compared to female flies (D.F. Duneau et al. 2017). Another study demonstrated that Tsf1 levels in the hemolymph are significantly higher in adult male flies (22 μM) compared to female flies (4.4 μM) (Weber et al. 2022). These results introduce the possibility of attributing differential disease outcomes to differences in iron levels between male and female flies. However, a direct link between iron sequestration and the sexually dimorphic survival of flies following infection has not yet been established. In this study we investigate whether iron sequestration is sexually dimorphic and if it contributes to sex differences in survival outcome.

In this study, we found evidence that the sexually dimorphic survival is linked to lower iron levels in male hemolymph caused by higher *Tsf1* expression in males compared to females. These differences are likely caused by the sexually dimorphic activation of the Toll-pathway. Altogether, we show that iron sequestration, a nutritional immune response, is sexually dimorphic and influences the response to infection between the sexes.

## Materials and Methods

### Drosophila stocks and rearing

The following *Drosophila* stocks used in this study were described previously: DrosDel *w*^*1118*^ iso; Oregon R; *Relish*^*E20*^ iso; *spz*^*RM7*^ iso; *Tsf1*^*JP94*^ iso; *c564-GAL4; UAS-Tsf1; UAS-CD8-GFP* (Iatsenko et al. 2020; Marra et al. 2021; Shaka et al. 2022; Arias-Rojas et al. 2023; Rubinić et al. 2025). The stocks were routinely maintained at 25°C with 12/12 h dark/light cycles on a standard cornmeal-agar medium: 3.72g agar, 35.28g cornmeal, 35.28g inactivated dried yeast, 16 ml of a 10% solution of methyl-paraben in 85% ethanol, 36 ml fruit juice, 2.9 ml 99% propionic acid for 600 ml. Fresh food was prepared weekly to avoid desiccation. Flies were flipped to new vials with fresh food every 3-4 days to grow new generations.

### Pathogen strains and survival experiments

In this study we used *Providencia alcalifaciens* DSM30120 that was obtained from the German Collection of Microorganisms and Cell Cultures (DSMZ) and *Providencia rettgeri* (Galac and Lazzaro 2011). The strains were grown in LB media (Invitrogen) overnight at 37°C in a shaking incubator. The culture was pelleted by centrifugation to remove the media and diluted to the desired optical density (OD600 = 0.25, 0.5, 1, 2, 3) with sterile PBS. If not otherwise indicated flies were infected with OD600 = 2. To infect flies, a 0.15 mm minutien pin (Fine Science Tools) mounted on a metal holder was dipped into the diluted bacterial solution and poked into the thorax of a CO2 anesthetized fly. Infected flies were maintained in vials (20 flies per vial) with food at 25°C overnight and moved to 29°C around 15-18h post infection. Surviving flies were counted at regular intervals. For the survival experiments that involved the prefeeding of flies with chemical compounds or sucrose, flies were fed for 24h with a mix of sucrose+5mM Ferric ammonium citrate (FAC) or 2.5% sucrose (control group). The respective solution was applied on top of a filter disc covering the fly food. Flies were flipped into fresh vials without the filter after infection.

### Quantification of pathogen load

For bacterial counts, flies were infected with *P. alcalifaciens* as described above and kept at 25°C. The number of bacteria was determined as follows at 6 h and 16 h post-infection: flies were surface sterilized in 70% ethanol, 3x for 5 seconds and then washed in sterile PBS 1x for 5 seconds. Flies (1 fly per replicate) were homogenized in 500 μl of sterile PBS for 30 s at 7200 rpm using a Precellys 24 instrument (Bertin Technologies, France). Serial 10-fold dilutions were made and plated on LB culture medium. The plates were left to dry and incubated overnight at 37°C. Colonies were counted and CFU were calculated as described previously (Hrdina et al. 2024).

### RT-qPCR

For quantification of messenger RNA, whole flies (n = 10) were collected at indicated time points. Total RNA was isolated using TRIzol Reagent (Invitrogen) and dissolved in Rnase-free water. 500 ng of total RNA was then reverse transcribed in 10 μL reactions using PrimeScript RT (TAKARA) and random hexamer primers. The qPCR was performed on a LightCycler 480 (Roche) using the the SYBR Select Master Mix from Applied Biosystems. RP49 was used as a housekeeping gene for normalization. Primer sequences were published previously (Hrdina et al. 2024).

### Hemolymph extraction and colorimetric iron measurement

To extract hemolymph, about 100 flies were anesthetized and placed on two 10 μm filter inside of an empty Mobicol spin column (MoBiTec). Glass beads were added on top of the flies and columns were centrifuged for 10 min at 4 °C, 5,000 rpm. The hemolymph was collected in 50 μL of Protease Inhibitor cocktail (Sigma, Catalog #11697498001, 1 tablet in 4ml PBS). Then each sample was diluted in a 1:10 ratio and the amount of protein was measured using the Pierce BCA Protein Assay Kit (Thermo Fisher Scientific) according to the manufacturer’s protocol. Iron was quantified using a colorimetric assay as was previously described in Xiao et al. 2019 (Xiao et al. 2019). Iron concentration in each sample was normalized to the total protein amount to standardize sample size differences and 120 μg were used as the sample in each assay. Protein samples (filled up to 50 μL with Protease Inhibitor Cocktail) were hydrolyzed with 11 μL of 32% Hydrochloric acid (VWR chemicals) under heating conditions (95 °C) for 20 min and centrifuged for 10 min at 20°C, 16000g. 18 μL of 75 mM Ascorbate (Sigma Aldrich) was added to 45 μL of supernatant followed by 18 μL of 10 mM Ferrozine (Sigma Aldrich) and 36 μL Ammonium Acetate (Chem-Lab NV). Absorbance was measured at 562 nm using an Infinite 200 Pro plate reader (Tecan). Quantification was performed using a standard curve generated with serial dilutions of a 10 mM FAC stock dilution.

### Quantification and statistical analysis

Data representation and statistical analysis were performed using the GraphPad Prism 10 software. Each experiment was repeated independently a minimum of three times (unless otherwise indicated), error bars represent the standard deviation (SD) of replicate experiments. The survival graphs show the cumulative survival analysed using the Cox-proportional hazard model. Other data was analyzed using Two-way ANOVA. Where multiple comparisons were necessary, appropriate Tukey or Sidak post hoc tests were applied.

## Results

### Sexual dimorphism in *Drosophila* resistance to *P. alcalifaciens* infection

To address the role of iron sequestration in sexually-dimorphic infection outcome, first we wanted to establish a suitable model. We decided to use *P. alcalifaciens* – a pathogen with a previously demonstrated higher virulence to female flies (D.F. Duneau et al. 2017). Importantly, Tsf1-mediated iron sequestration is an effective host defense mechanism against *P. alcalifaciens* (Shaka et al. 2022). Hence, this pathogen can be used to investigate the link between sexual dimorphism in iron sequestration and infection susceptibility. We confirmed increased susceptibility of female flies to *P. alcalifaciens* infection using w1118 iso and Oregon R flies (Fig. 1a and Supplementary Figure 1a). This phenotype was reproducible across different pathogen doses (Supplementary Figure 1b-e) and was observed with another *Providencia* sp -*P. rettgeri* (Fig. 1b). Given that mating and reproduction have been shown to suppress immune responses in *D. melanogaster* females (Schwenke et al. 2016; Gordon et al. 2025), we tested whether increased susceptibility of females is a consequence of immunosuppressive effect of reproduction. As shown in Fig 1c, sex differences in survival were still observed in virgin flies, indicating that the higher susceptibility of female flies is not due to a higher reproduction-immunity tradeoff than in males. Consistent with an increased susceptibility to infection, female flies contained higher pathogen loads than male flies (Fig. 1d). While it was statistically significant only at 16h post-infection, at 6h the mean CFU value was 5.76 times higher in females compared to males, suggesting that the pathogen proliferates faster in female hosts.

**Figure 1.**
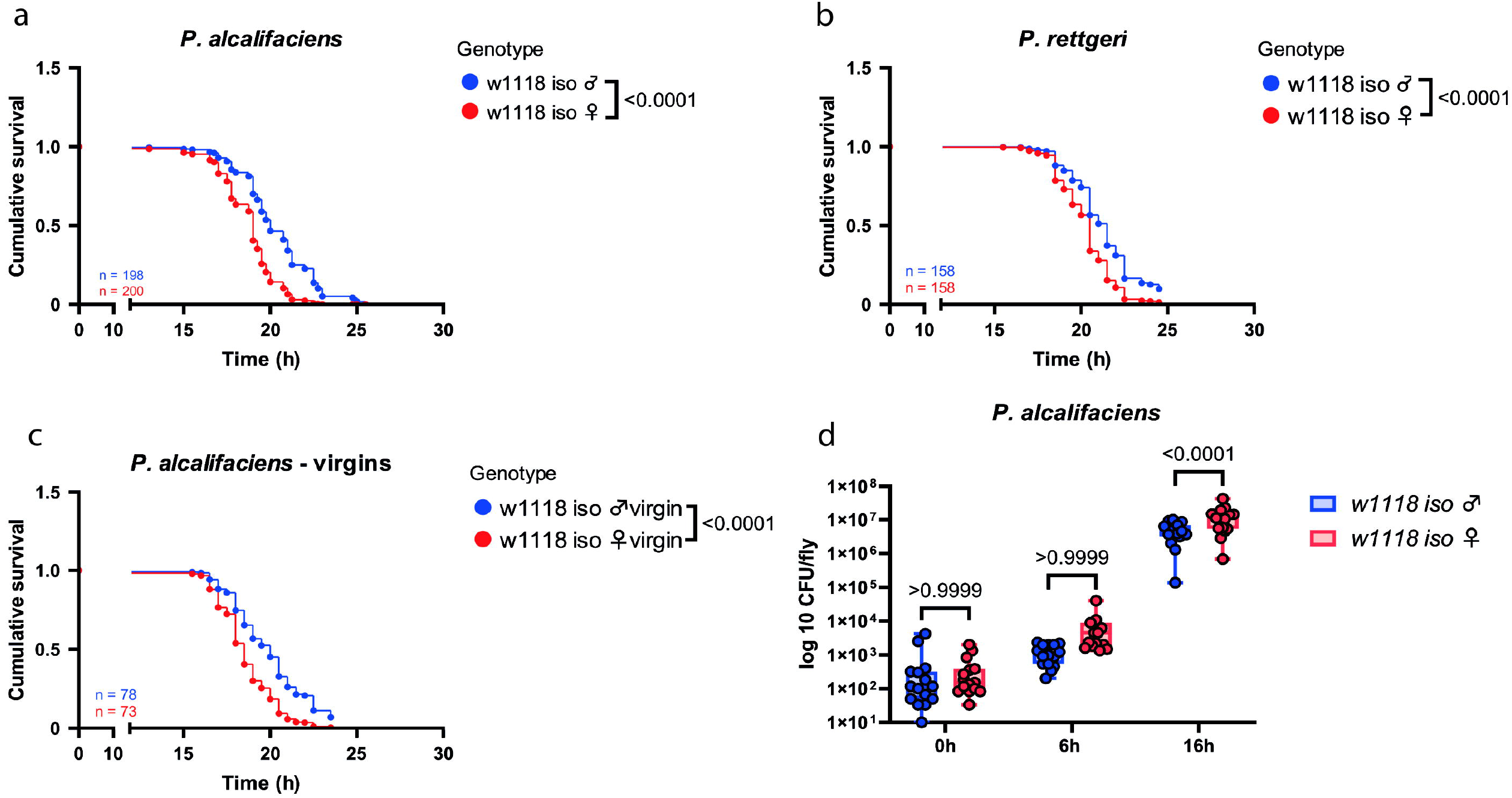
Male flies are more resistant to infection with *Providencia sp* than female flies. **(a-c)** Cumulative survival of mated **(a)** or virgin **(c)** adult flies infected with *P. alcalifaciens* and mated adult flies infected with *P. rettgeri* **(b)**. n indicates the total number of flies used in the experiments per genotype. **(d)** *P. alcalifaciens* load at 0h, 6h and 16h post infection. For CFU counts, one dot represents the pathogen load of one infected fly. Results are shown as mean and SD of 3 independent experiments.

### Sex-biased *Tsf1* expression leads to sexually-dimorphic iron sequestration

To investigate the role of Tsf1 in the observed sex differences in infection susceptibility, we first compared *Tsf1* expression between males and females using *Tsf1* expression data extracted from Flyatlas (Leader et al. 2018). As shown in Fig 2a, *Tsf1* expression was significantly higher in males than in females in the whole body and in most of the tissues tested. Using qPCR, we confirmed a significantly higher basal expression level of *Tsf1* in males. While *P. alcalifaciens* infection induced *Tsf1* expression in both sexes, the expression level remained significantly higher in male flies at both time points post infection (Fig. 2b).

**Figure 2.**
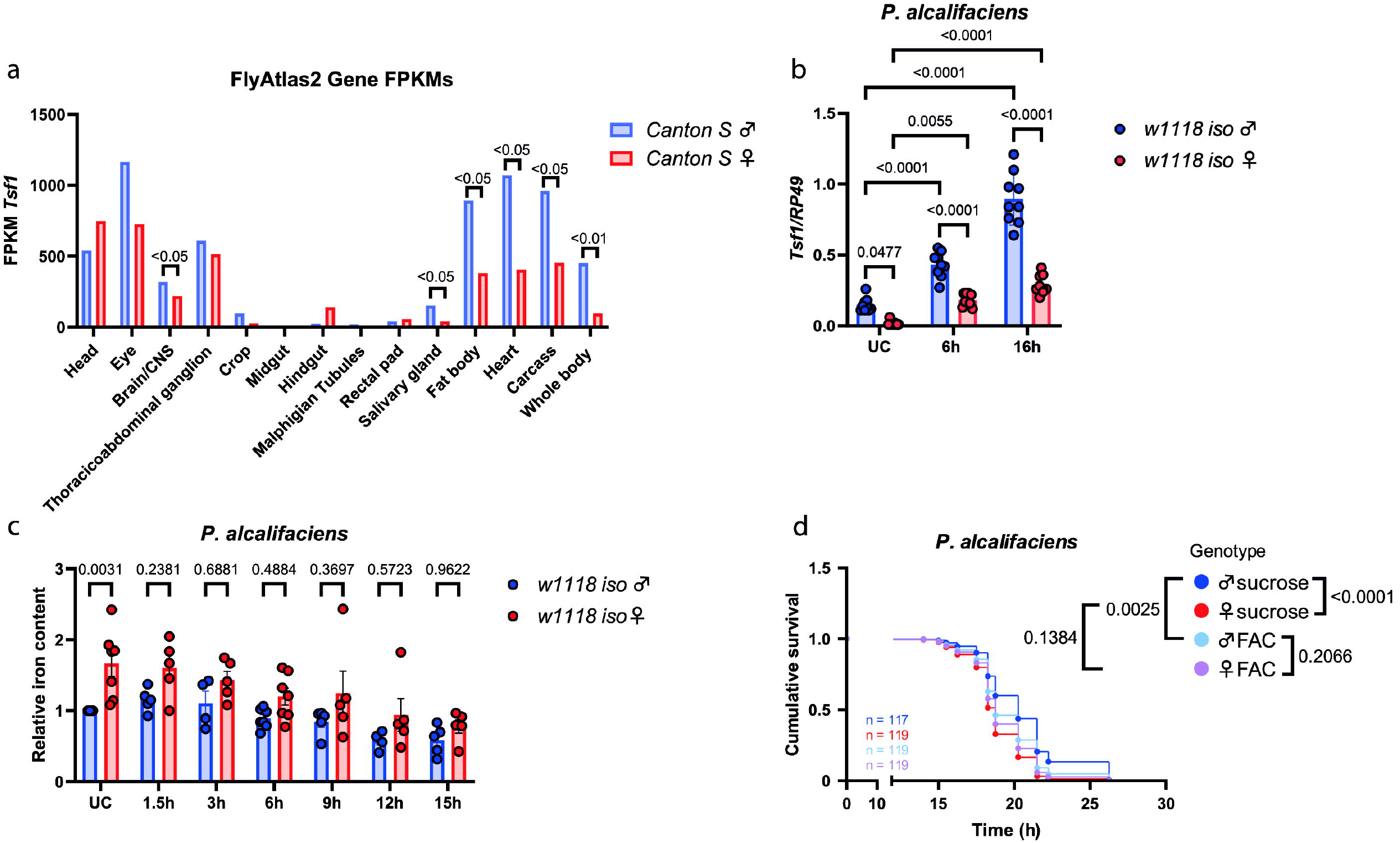
Male flies show higher Tsf1 levels and lower basal iron content in the hemolymph than female flies. **(a)** Expression data of *Tsf1* in whole flies and specific tissues taken from FlyAtlas 2. **(b)** RT-qPCR measuring *Tsf1* expression in unchallenged (UC) or flies infected with *P. alcalifaciens* 6 and 16 hours post infection, respectively. One dot represents expression levels from 10 pooled flies. The mean and SD of three independent experiments are shown. **(c)** Relative iron levels in the hemolymph of unchallenged (UC) flies or after infection with *P. alcalifaciens* after indicated timepoints. The iron content was measured using the ferrozine assay. One dot represents iron levels of a pool of 75+ flies. Unchallenged male flies were set to 1 and all other data show the relative amount of this value. The mean and SD of at least three independent experiments are shown. **(d)** Cumulative survival of *w1118 iso* flies that were previously fed with FAC or sucrose as a control for 24h. n indicates the total number of flies used in the experiments per genotype.

Given the prominent role of Tsf1 in infection-induced iron sequestration, we tested whether sex-biased survival correlates with sex differences in iron sequestration. We measured the iron content in the hemolymph of uninfected and infected with *P. alcalifaciens* male and female flies using a ferrozine assay. As shown in Fig 2c, iron levels were significantly higher in female flies than in male flies under uninfected conditions. We have not detected any significant differences in iron levels between males and females at various timepoints after *P. alcalifaciens* infection, although there was a trend towards higher iron load in female flies (Fig 2c). These results suggest that basal differences between sexes in iron content prior to infection might contribute to sexual dimorphism in survival. To test this possibility, we prefed flies with Ferric ammonium citrate (FAC) prior to infection to saturate iron load. FAC feeding increased male susceptibility to *P. alcalifaciens* infection, while there was no significant effect of FAC on female survival (Fig 2d). Importantly, in contrast to control sucrose-fed flies, FAC-fed flies exhibited no sex differences in survival to *P. alcalifaciens* infection (Fig 2d). Hence, increasing iron levels through FAC feeding makes males more susceptible to infection and eliminates sex differences in survival.

### Tsf1 mediates sexual dimorphism in survival and iron sequestration

To investigate whether Tsf1 mediates sex differences in infection susceptibility and iron content, we assessed the survival of a *Tsf1* mutant to *P. alcalifaciens* infection. As expected from our previous work (Shaka et al. 2022), *Tsf1* mutants were more susceptible to *P. alcalifaciens* infection (Fig. 3a). Notably, in contrast to wild-type flies, *Tsf1* mutants did not show significant differences between males and females in survival (Fig. 3a). Similar results were observed with *P. rettgeri* (Fig. 3b). Therefore, Tsf1 is an important mediator of sexual dimorphism in survival to some pathogens. Consistent with survival, we detected higher pathogen loads in *Tsf1* mutant males and females as compared to wild-type (Fig. 3c). As anticipated, *P. alcalifaciens* reached higher titer in wild-type females than in males. Interestingly, sex dimorphism in *P. alcalifaciens* load was still present in the *Tsf1* mutant, which showed no sex differences in survival (Fig. 3c). Thus, Tsf1 might play a sexually dimorphic role in host tolerance to *P. alcalifaciens* infection.

**Figure 3.**
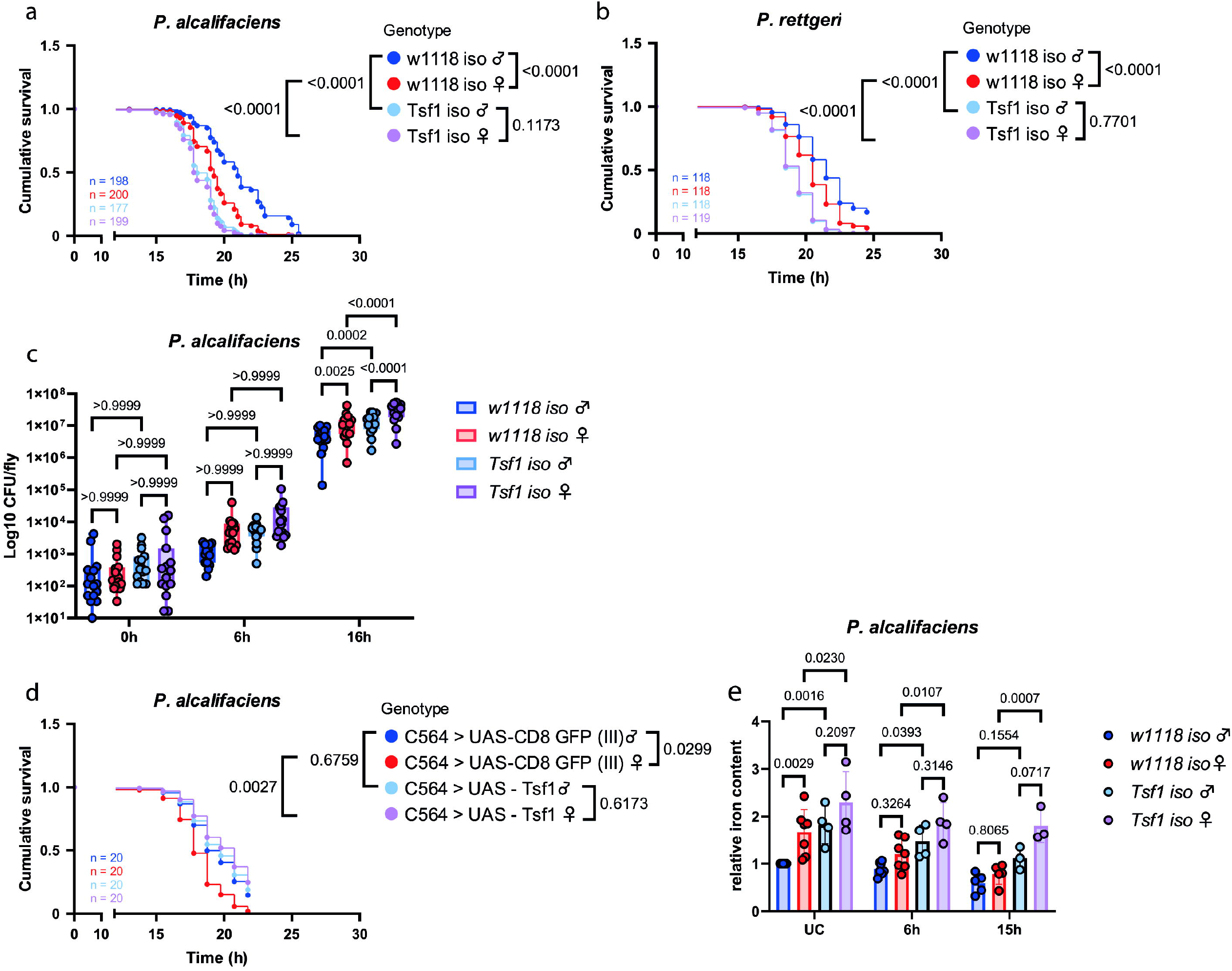
Differences in *Tsf1* expression cause the sexual dimorphism in survival and iron content. **(a-b)** Cumulative survival of adult flies infected with *P. alcalifaciens* **(a)** and *P. rettgeri* **(b)**, respectively. n indicates the total number of flies used in the experiments per genotype. **(c)** *P. alcalifaciens* load at 0h, 6h and 16h post infection. For CFU counts, one dot represents the pathogen load of one infected fly. Results are shown as mean and SD of 3 independent experiments. **(d)** Cumulative survival of flies overexpressing Tsf1 (C564 > UAS-Tsf1) or GFP as a negative control (C564 > UAS-CD8 GFP) in the fat body. n indicates the total number of flies used in the experiments per genotype. **(e)** Relative iron levels in the hemolymph of unchallenged (UC) flies or after infection with *P. alcalifaciens* after 6h and 15h. The iron content was measured using the ferrozine assay. One dot represents iron levels of a pool of 75+ flies. Unchallenged *w1118 iso* male flies were set to 1 and all other data show the relative amount of this value. The mean and SD of at least three independent experiments are shown.

To reinforce the role of Tsf1 in sexual dimorphism, we measured the survival of flies with *Tsf1* overexpression in the fat body. In contrast to the control line overexpressing GFP, the *Tsf1* overexpressing line showed no significant differences between sexes in survival (Fig. 3d). This loss of sex differences could be attributed to an increased survival of female flies with *Tsf1* overexpression, as such overexpression had no significant effect on males’ survival (Fig. 3d).

Finally, measuring iron levels revealed that *Tsf1* mutant flies contained more iron in the hemolymph compared to wild-type flies (Fig. 3e). However, in contrast to wild-type flies which showed sexual dimorphism in basal iron load, no significant differences between sexes were identified in the *Tsf1* mutant flies (Fig. 3e), suggesting that Tsf1 mediates sexual dimorphism in the hemolymph iron content.

### Sex differences in the Toll pathway activation determine sexual dimorphism in *Tsf1* expression and fly survival

Next, we wanted to identify the signalling pathway controlling the sex-biased *Tsf1* expression. Given that *Tsf1* expression can be controlled by the Imd or Toll pathway depending on the pathogen (Iatsenko et al. 2020), we investigated whether any of these pathways contribute to sexually-dimorphic expression of *Tsf1*.

To test the role of the Imd pathway, we measured *Tsf1* expression in a *Relish* mutant. As shown in Fig 4a, *Tsf1*expression after infection was significantly lower in *Relish* mutant as compared to wild-type flies in both sexes. However, the sexual dimorphism in *Tsf1* expression was still observed in *Relish* mutant flies, as illustrated by the significant differences in the expression between males and females after infection. These results demonstrate, that while Imd pathway is required for the full *Tsf1* induction after infection, it does not mediate sex differences in *Tsf1* expression. Next, we investigated the contribution of the Toll pathway using *spz* mutant and found that in *spz* mutant sex differences in *Tsf1* expression were not present (Fig 4b). However, *Tsf1* expression was induced in *spz* mutant by infection to the same degree as in wild-type flies (Fig 4b). Therefore, the Toll pathway is not required for *P. alcalifaciens* - induced *Tsf1* expression but it mediates sex differences in *Tsf1* expression under basal and infection conditions. Consistent with the expression data, we found that *Relish* mutants exhibited sex differences in survival with females being more susceptible (Fig 4c). In *spz* mutants, we did not find significant differences in survival between males and females (Fig 4d). Hence, the Toll pathway mediates sex differences in *Tsf1* expression and susceptibility to infection.

**Figure 4.**
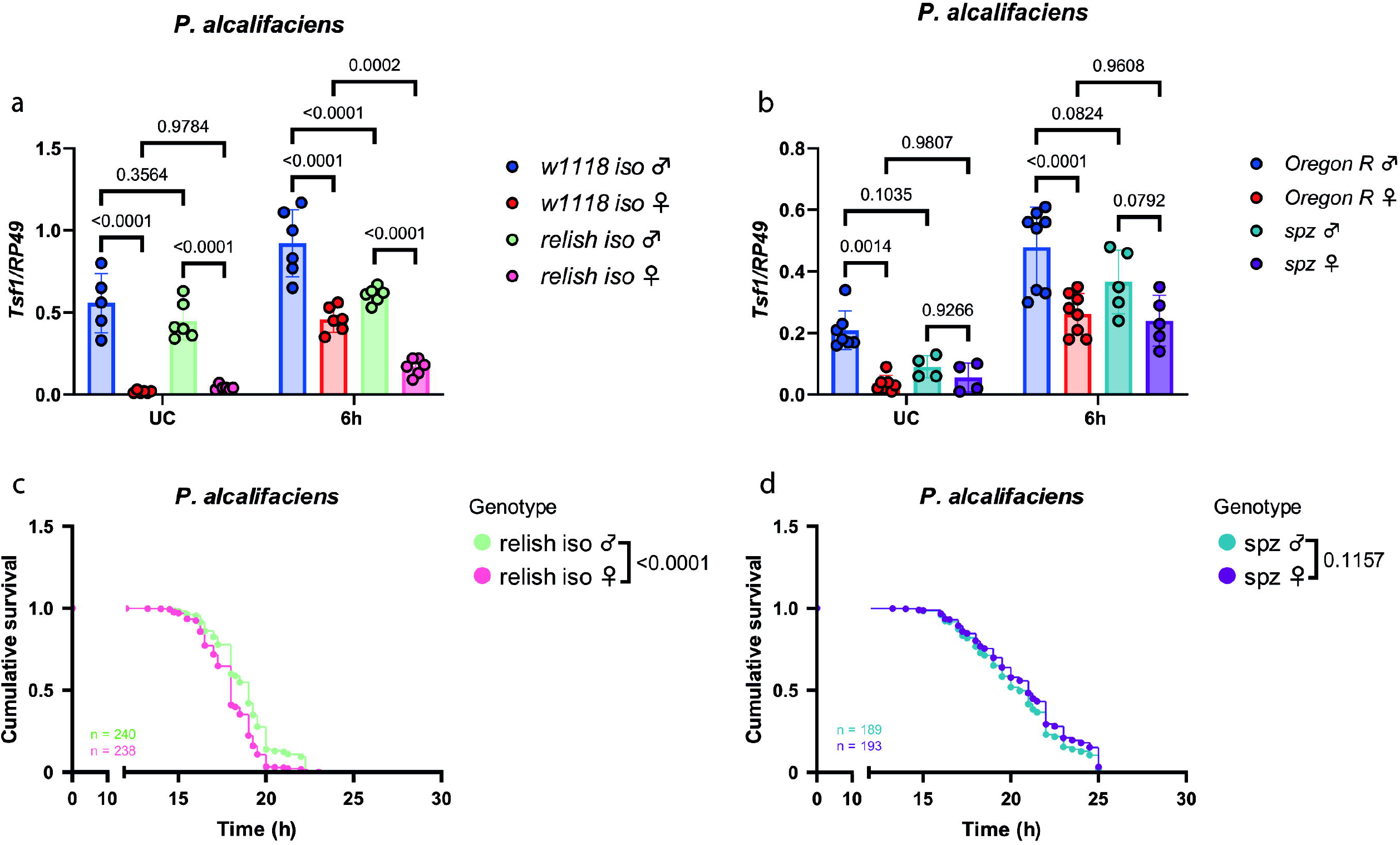
Differences in Toll pathway activation induce the sexually dimorphic *Tsf1* expression and fly survival. **(a-b)** RT-qPCR measuring *Tsf1* expression in IMD-(*relish iso*) **(a)** and Toll-(*spz*) **(b)** pathway mutants. Data show unchallenged (UC) or flies infected with *P. alcalifaciens* 6 and 16 hours post infection, respectively. One dot represents expression levels from 8-10 pooled flies. The mean and SD of at least two independent experiments are shown. **(c-d)** Cumulative survival of immune mutants *relish iso* (IMD-pathway) **(c)** and *spz* (Toll-pathway) **(d)**. n indicates the total number of flies used in the experiments per genotype.

## Discussion

Across taxa, understanding the differences between males and females in infection outcomes remains a significant unresolved question. In this study, we showed that sex differences in nutritional immunity, specifically iron sequestration, contribute to the sexually dimorphic susceptibility of fruit flies to *P. alcalifaciens* infection. Specifically, male flies exhibit elevated expression of a key iron transporter – Tsf1 compared to females. This resulted in a lower amount of iron in hemolymph of males. Given that iron is an essential element for bacterial growth, lower iron content in males leads to reduced pathogen growth and significantly longer survival of male flies. Importantly, while *Tsf1* expression was consistently higher in males before and during infection, sex differences in iron content were significant only under basal conditions. Given that initial phase of infection predicts the probability of survival in *Drosophila* (D. Duneau et al. 2017), basal differences between males and females in the amount of iron that is available to pathogens are likely to be sufficient to determine the sex-biased infection outcome. Indeed, we detected higher pathogen load in female flies, which likely stems from an increased availability of iron.

Further support for the link between iron availability and susceptibility to *P. alcalifaciens* infection comes from FAC feeding and *Tsf1* overexpression experiments. The increase of iron load by FAC feeding significantly increased susceptibility of males but not females to *P. alcalifaciens* infection, while decreasing iron load by *Tsf1* overexpression increased female but not male survival after infection. Such sex-biased effects of these treatments could be attributed to the pre-existing sex differences in iron content. Since the basal iron load in females is higher compared to males, further increase by FAC feeding, if any, has no consequences for their survival. While with *Tsf1* overexpression the situation is opposite – further decrease of already low iron content is not relevant for male survival. Together, these experiments confirm a sexual dimorphism in iron metabolism and its role in susceptibility to infection.

The sex differences in iron sequestration that we identified could potentially explain other sexually dimorphic traits in *Drosophila*. Since iron is a redox-active metal, its excess can promote the generation of ROS. Hence, elevated iron levels in the hemolymph of female flies might be the reason for their increased ROS levels and susceptibility to oxidative stress (Albrecht et al. 2011; Rubinić et al. 2025). On the other hand, increased ROS levels might provide protection to female flies and explain their increased resistance against certain pathogens, like *S. aureus*, which is believed to be susceptible to immune-induced ROS (Dudzic et al. 2019). Since certain pathogens, like the microsporidian *Nosema ceranae*, and endosymbionts, such as *Spiroplasma*, utilize iron complexed with transferrin as an iron source within the host environment (Rodríguez-García et al. 2021; Marra et al. 2021), increased availability of Tsf1 in male flies might render them more susceptible to such pathogens.

We found that the Toll pathway controls basal and infection-induced differences between males and females in *Tsf1* expression. Given a previously demonstrated elevated activity of the Toll pathway in male flies (D.F. Duneau et al. 2017), elevated *Tsf1* expression and iron sequestration in males are a likely consequence of higher Toll pathway activity. Hence, we identified Tsf1 as one of the Toll pathway-regulated immune effectors involved in immune dimorphism, further deepening our understanding of the mechanisms underlying sex-biased susceptibility to infection. Given that the Toll pathway controls the expression of multiple immune effectors, it is possible that besides Tsf1, additional Toll-regulated immune effectors contribute to sex differences in the susceptibility to various infections. However, in case of *P. alcalifaciens*, Tsf1 is a key mediator of sexual dimorphism as sex differences in survival were abrogated in *Tsf1*-deficient flies. Notably, while the differences in iron levels in the *Tsf1* mutant males and females were not statistically significant, there was still a trend towards higher iron amount in *Tsf1* females. This indicates that, in addition to Tsf1, there are likely to be additional iron transporters, which remain to be identified, that might mediate sex differences in the hemolymph iron content.

The important remaining question is what determines sexual dimorphism in the Toll pathway. Given the established interactions between immunity and hormones in *Drosophila* (Regan et al. 2013; Schwenke et al. 2016), hormonal differences between females and males are likely contributors to differential Toll pathway activation. For instance, females produce juvenile hormone (JH) which is crucial for reproduction. JH suppresses the immune response with a stronger influence on the Toll pathway than on the Imd pathway (Flatt et al. 2008; Schwenke and Lazzaro 2017). Therefore, the female-biased production of JH has a cost on their immune system, leading to sexual dimorphism in the Toll pathway activity and response to certain infections (D.F. Duneau et al. 2017).

Consistent with our previous work (Shaka et al. 2022), we found that *Tsf1* mutants of both sexes were more susceptible to *P. alcalifaciens* infection and exhibited a higher pathogen load than wild-type flies. Therefore, *Tsf1* mutants have impaired resistance mechanisms compared to wild type flies or a reduced ability to clear the pathogen. However, we noticed that *Tsf1* females, while having the same survival rate as *Tsf1* males, carried a significantly higher pathogen load at 16h post-infection. Therefore, *Tsf1* mutant females are more tolerant than males to *P. alcalifaciens* infection as they can sustain higher pathogen burden and associated pathology without apparent consequences for survival. These findings uncover a previously unrecognized role of Tsf1 in sex differences in host tolerance to infection. Given an increasing interest in understanding sex differences in tolerance to infection (Howick and Lazzaro 2017; Vincent and Dionne 2021), further dissection of the mechanisms behind increased tolerance of *Tsf1* females to *P. alcalifaciens* will provide valuable insights into the biological processes that contribute to sex-specific immune tolerance.

## Data availability

The authors affirm that all data necessary for confirming the conclusions of this article are represented fully within the article and its tables and figures.

## Acknowledgements

We are grateful to the Bloomington Drosophila Stock Center (NIH P40OD018537) for fly stocks. We thank Dagmar Frahm for technical assistance.

## Author contributions

AH – investigation, formal analysis, data curation, methodology, visualization, writing – original draft preparation, writing – review and editing; II – conceptualization, funding acquisition, project administration, supervision, resources, writing – original draft preparation, writing – review and editing.

