## Supplementary figures and images for "Sex Bias in Iron Sequestration by Transferrin 1 Modulates Sexually-Dimorphic Infection Outcomes in *Drosophila melanogaster*"

### Supplemental Figure 1

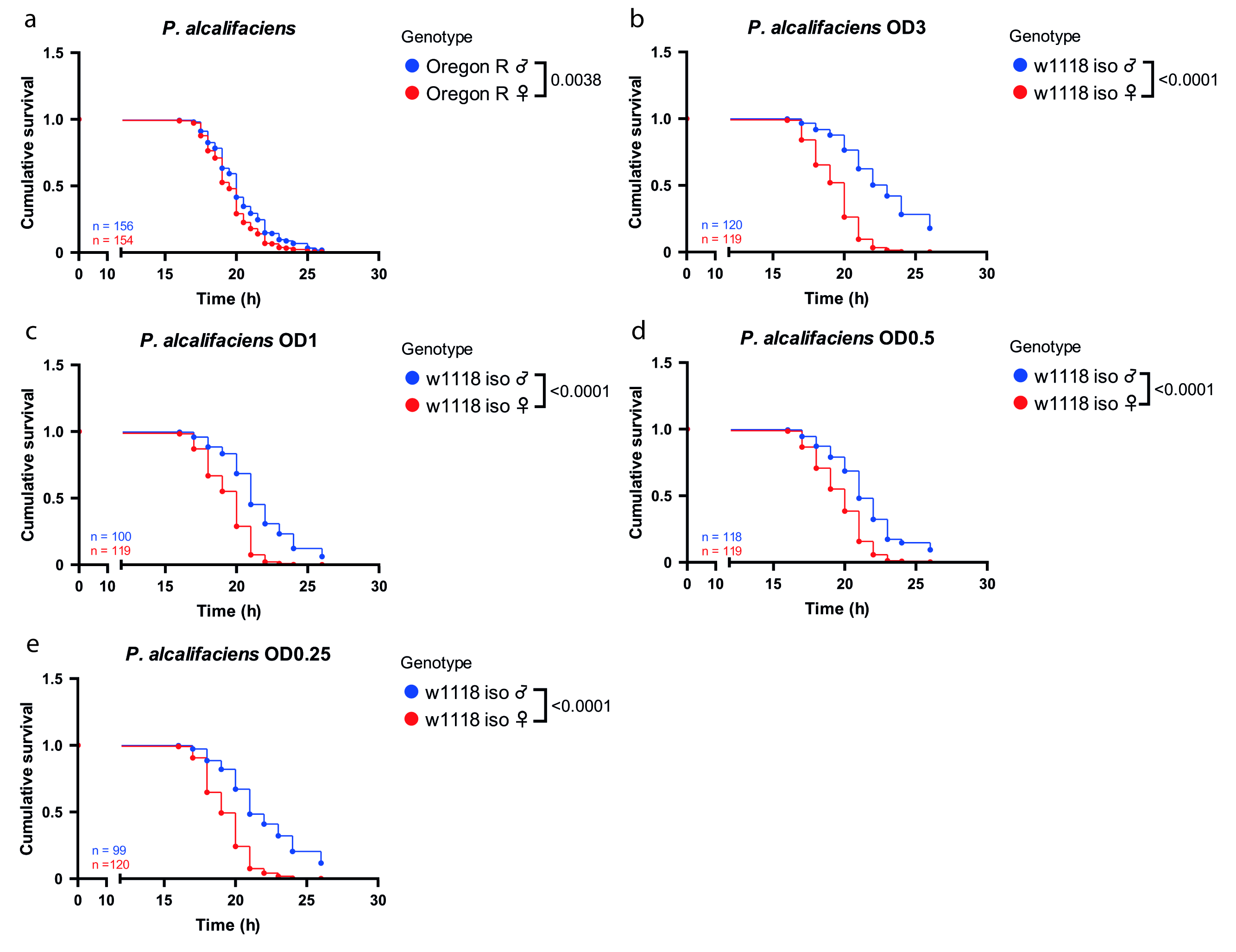
